# Unbiased discovery of natural sequence variants that influence fungal virulence

**DOI:** 10.1101/2023.04.23.537984

**Authors:** Daniel Paiva Agustinho, Holly Leanne Brown, Guohua Chen, Michael Richard Brent, Tamara Lea Doering

**Author notes:** Human Genome Sequencing Center, Baylor College of Medicine, Houston, TX, 77030. Pfizer, Inc., Chesterfield, MO, 63107.

## Abstract

Isolates of *Cryptococcus neoformans*, a fungal pathogen that kills over 120,000 people each year, differ from a 19-megabase reference genome at a few thousand up to almost a million DNA sequence positions. We used bulked segregant analysis and association analysis, genetic methods that require no prior knowledge of sequence function, to address the key question of which naturally occurring sequence variants influence fungal virulence. We identified a region containing such variants, prioritized them, and engineered strains to test our findings in a mouse model of infection. At one locus we identified a 4-nt variant in the *PDE2* gene, which severely truncates its phosphodiesterase product and significantly alters virulence. Our studies demonstrate a powerful and unbiased strategy for identifying key genomic regions in the absence of prior information, suggest revisions to current assumptions about cAMP levels and about common laboratory strains, and provide significant sequence and strain resources to the community.

## INTRODUCTION

The pathogenic fungus *Cryptococcus neoformans* causes lethal meningoencephalitis that kills 112,000 people worldwide each year (1). Researchers in this field have collected thousands of *C. neoformans* isolates, which have been used to elucidate *C. neoformans* evolution (2-14) and in efforts to correlate disease outcome with *in vitro* measures such as virulence factor production or fungal growth (5,15-19). It is clear that distinct strain lineages are associated with varied clinical outcomes (5,10,14,20-24). Furthermore, natural genomic sequence variation has been associated with levels of virulence, both in human infection and in mouse studies (21,22,25,26). However, the identification of specific sequence variants that causally influence the virulence of clinical strains has remained a considerable challenge. (Here we use variants to refer to nucleotide substitutions or indels shorter than 50 bp.)

We have developed and validated an unbiased, genetic strategy to identify naturally occurring sequence variants that impact *C. neoformans* virulence. Our whole-genome approach can potentially reveal key variants in novel genes, directing research attention to their products. It can also highlight specific variants in genes that have already been implicated in virulence, which can lead to mechanistic understanding of the encoded proteins. Additionally, and in contrast to studies based on gene deletion, our strategy can identify critical variants in essential genes, regulatory sequences, and non-annotated or mis-annotated regions of the genome.

Valuable information relating fungal genotype to disease outcome has been derived from strains isolated from patients with cryptococcosis and their accompanying clinical records. One challenge of such studies, however, is that complex host factors contribute to outcome, including patient genotype, known and unknown comorbidities, treatment, and healthcare setting (27). This complexity limits the power of these analyses. To circumvent this, we used mouse models of infection to assess strains derived from clinical isolates. This approach has been validated in the literature, which shows a strong correlation between mortality in humans and mice infected with the same *C. neoformans* strain (20). Another challenge for our plans to exploit genetic analysis is the large haplotype blocks observed in clinical isolates, a result of limited recombination in the wild (6,14). To address this, we have taken advantage of the sexual cycle of *C. neoformans* (28,29), which allows us to generate recombinant progeny for study.

We applied genetic approaches to analyze recombinant progeny derived from a cross between a well characterized and highly virulent laboratory strain (KN99 (30)) and a clinical strain that exhibits low virulence in our mouse model (C8 (31)). Excitingly, our unbiased approach efficiently identified individual causal variants responsible for virulence differences among *C. neoformans* isolates. We identified and experimentally validated sequence variants that both increase and decrease virulence, showing that even relatively less pathogenic strains harbor variants that increase virulence. We also showed that the phosphodiesterase Pde2 lacks activity in all *Cryptococcus neoformans* laboratory strains, significantly reducing their virulence.

## RESULTS

The first step in our strategy was to select clinical isolates for genetic crosses. For one parent, we chose *C. neoformans* strain KN99**a**, which is the congenic partner of KN99α (30), a reference genome strain for *C. neoformans* (32). The KN99 strains, which are derived from the clinical isolate H99 (33), are highly virulent in animal models and mate robustly (30). For the second parent we wanted a clinical isolate that differed significantly from KN99 in terms of genome sequence and virulence, but also mated well, which is not typical of these isolates. To identify such a strain among the 73 clinical isolates that our laboratory had on hand, we first compared their genome sequences (either obtained online or sequenced in-house; see Methods and Supplemental Table 1, sheet A) to that of KN99α. This analysis yielded a total of 1,072,542 distinct genomic variants relative to KN99.

Even when patient data is available, it is challenging to directly compare the virulence of clinical isolates because of the confounding host factors mentioned above. For this reason, we used an animal model of cryptococcal infection to compare clinical strains to the laboratory strain KN99. To do this, we infected mice intranasally (to mimic the common route of infection in humans) and assessed colony forming units (CFU) in the lung nine days after infection as a proxy for virulence. We observed a 14,000-fold range in lung burden between clinical isolates, with strains that were both more and less virulent than KN99**a** (Figure 1, top).

**Figure 1.**
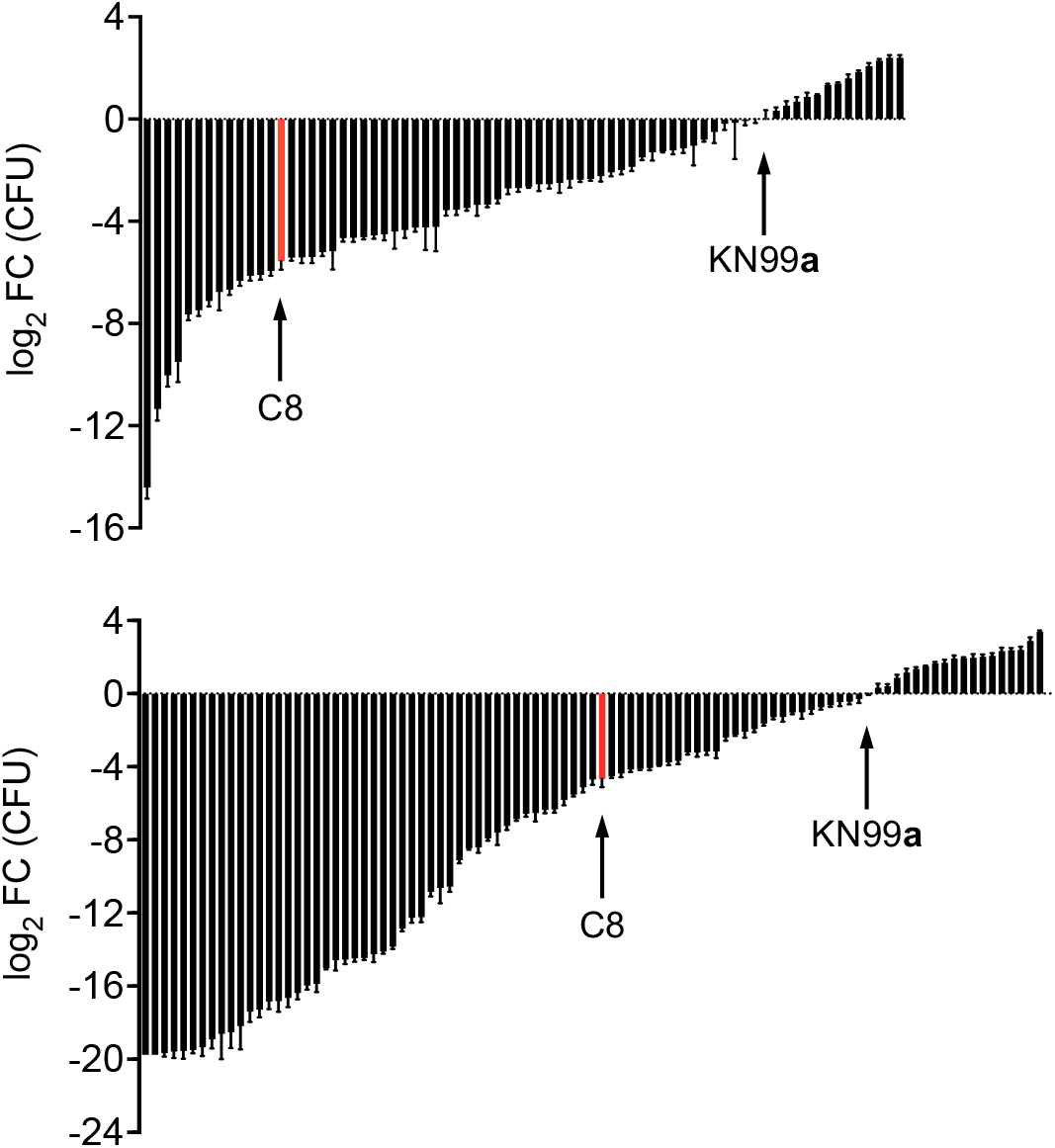
Individual C57BL/6J mice were intranasally inoculated with 12,500 cryptococcal cells of KN99**a**, C8 (orange bar), or other strains of interest. Lung burden was measured 9 days post-infection by colony forming units (CFU), which are plotted for each strain relative to the value for KN99**a**. *Top*, 73 clinical isolates. *Bottom*, 93 recombinants derived by crossing KN99**a** and C8.

Almost all of the clinical strains we examined, like most cryptococcal isolates, are mating type alpha (34). To evaluate their ability to produce mating structures, we crossed them to KN99**a** on V8 medium (35). Of the 73 strains, 34 exhibited some level of filamentation under our standard conditions (see Methods). We collected spores from four of these crosses by microdissection and sequenced at least five progeny strains from each. Two of the crosses produced progeny nearly identical to one of the parents (>99% of SNPs in common), possibly generated by the recently described process of pseudo- sexual reproduction (36). The other two generated the recombinant progeny expected from sexual reproduction; an example is depicted in Figure 2.

**Figure 2.**
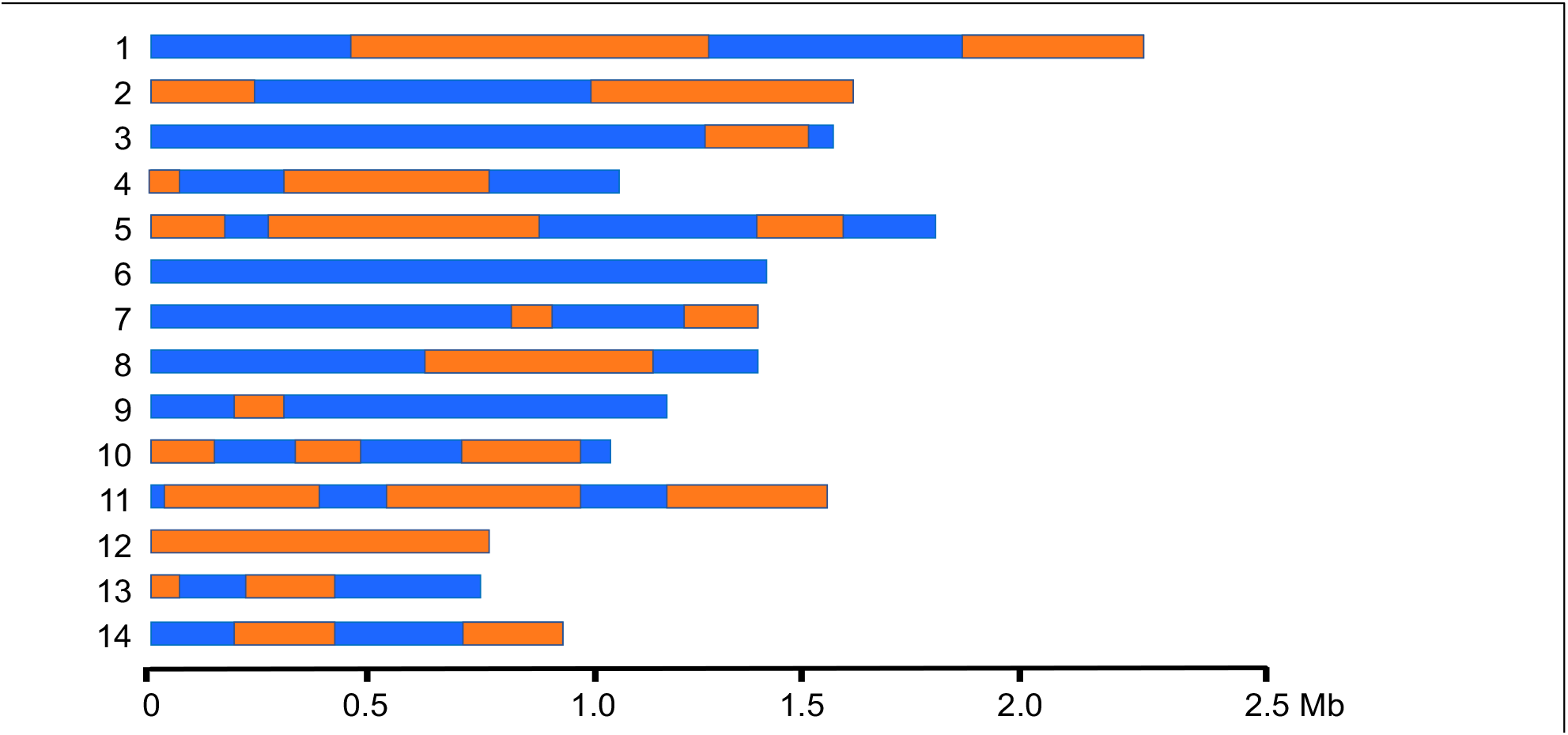
Representation of the whole genome sequence of one recombinant strain derived from a cross between C8 and KN99**a**. Each bar represents one of the 14 chromosomes of *C. neoformans*. Orange, haplotypes derived from C8; blue, haplotypes derived from KN99**a**.

We selected clinical strain C8 (31) (genome sequence accession SRX189616) for our studies, based on its virulence, mating characteristics, and production of recombinant progeny when crossed to KN99**a** (Figure 1). This strain, isolated from the cerebrospinal fluid of an HIV+ cryptococcosis patient in the United States (31), has 48,934 genomic variants (45,343 SNVs and 3591 indels) compared to KN99α, which are distributed throughout the genome. Notably, its virulence in our mouse model is ∼100 times lower than that of KN99**a** (Figure 1, top panel). We crossed these two strains and used microdissection to isolate 138 of the resulting spores, which were then cultured and stored as frozen stocks (see Methods and Supplemental Table 1, sheet B).

To identify sequence variants that would explain the virulence difference we observed between the KN99 and C8 parent strains, we took two distinct approaches: Bulked Segregant Analysis (BSA) and Association Analysis (AA). In BSA, which was first developed as a tool for plant genetic mapping (37), segregants from a single cross are divided based on a specific characteristic of interest and the resulting populations are subjected to molecular analysis. In our adaptation of this strategy, we collected populations of recombinants based on growth, either in rich laboratory medium or in the mouse lung after intranasal infection, and analyzed them by DNA sequencing. Our rationale was that alleles that are beneficial in either growth condition will increase in frequency, those that are deleterious will decrease, and those that are neutral will drift randomly. We postulated that alleles that are significantly enriched when cells are grown in the mouse lung are likely to be virulence-enhancing, while those that are significantly depleted are likely virulence-reducing. To identify such alleles, we randomly selected 100 of the C8 × KN99 progeny and combined them into five pools of 40 strains each, such that each strain was present in two distinct pools. Each pool was prepared in duplicate, with equal cell numbers of each strain. From these ten samples, aliquots were reserved as starting material; used to inoculate a culture of rich medium (YPD); and used to infect one mouse intranasally. We extracted DNA for sequencing from the starting material (inoculum), cells recovered from the YPD culture after 20 hours of growth at 30°C, and cells isolated from mouse lungs after 9 days of infection. To ensure that we could detect small changes in allele frequencies, all samples were sequenced to a minimum of 100-fold coverage by 2 × 150 bp, paired-end reads (Supplemental Table 1, sheet C).

Next, we evaluated each variant site in the genome for evidence of positive or negative selection by growth in YPD or mouse lung. To do this we compared each allele’s frequencies after growth to the frequency in the inoculum and calculated g’ statistics (38). We then summed the read counts across all pools from each condition for each allele and did the same frequency calculation. The top panel of Figure 3 shows the data for chromosome 2, with changes in allele frequency towards C8 arbitrarily assigned as positive and changes towards KN99 shown as negative. Results from the YPD samples (plotted in black) show little change in allele frequency from the starting material, with data points close to the x-axis all along the chromosome; this pattern was maintained throughout the genome (Supplemental Figure 1). In contrast, we observed large changes in allele frequency along this chromosome for samples that were isolated from the lungs of infected animals (plotted in red).

**Figure 3.**
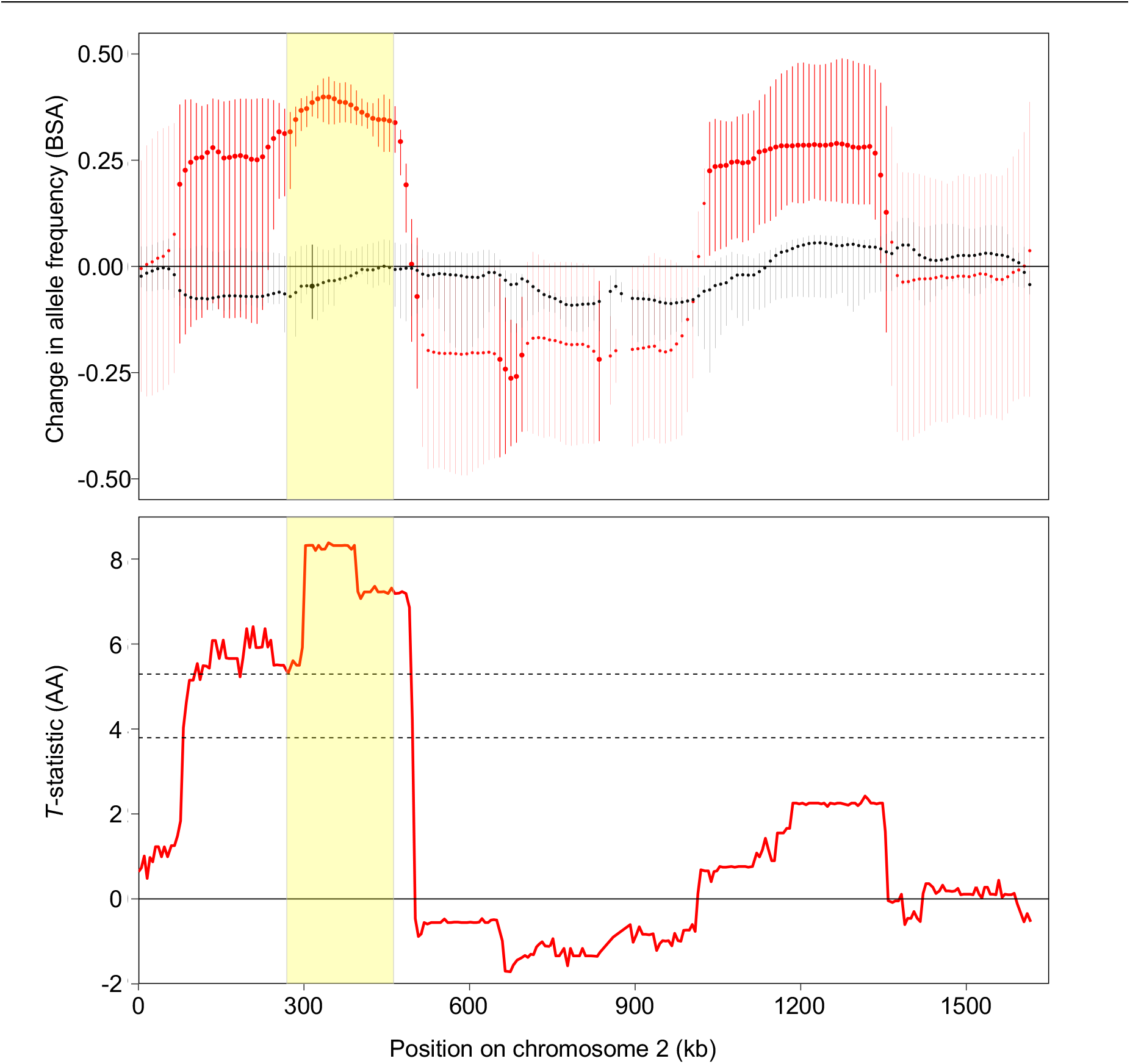
Distinct methods implicate the same genomic region in virulence. *Top panel*, changes in allele frequency for experimental samples compared to initial pools, with positive and negative values arbitrarily assigned to changes in the direction of C8 and KN99, respectively. Values are plotted for 5-kb windows of Chromosome 2, smoothed as described in the Methods. Symbols, mean value for all pools; vertical lines, range of individual pool values; black, YPD samples; red, mouse lung samples; yellow shaded region, IR-1 (see text). Larger symbols and darker lines indicate regions of statistical significance, as defined in the text. *Bottom panel*, association analysis results for all variants in Chromosome 2. The red line shows the t-statistic comparing the virulence (measured as log_2_ fold-change of CFU compared to KN99) of strains with each parental allele at each variable site along the chromosome. Dashed lines, P-value thresholds for 0.001 (top line) and 0.01 (bottom line). P-value calculations are detailed in the Methods.

We examined the BSA data for regions of the genome where changes in allele frequency were (a) statistically significant (false discovery rate < 0.05) for lung samples but not YPD samples and (b) varied in the same direction for the mean of the pools and for each individual pool. One such region, which shows close agreement between individual pools, occurs on chromosome 2 between positions 283,000 and 467,000 (highlighted in yellow on Figure 3, top panel). We termed this Implicated Region 1 (IR-1). It shows an increase in C8 allele frequencies in lung samples (red symbols), suggesting that one or more alleles in this region confers a growth advantage in this host environment. Changes in IR-1 allele frequencies for the pools grown in YPD (black symbols) were not significant, suggesting that the alleles in this region are neither beneficial nor deleterious for growth in rich medium.

We next used a completely different assay and statistical methodology to examine the same group of 100 progeny strains for variants important for virulence, by asking whether there was a statistical association between the presence of specific alleles and virulence in mice when strains were tested individually. To perform this association analysis (AA), we first tested each recombinant strain alone in mice (Figure 1, bottom panel), using lung burden (log_2_ fold-change in CFU compared to KN99) as a surrogate for virulence. Notably, the range of virulence of the recombinants greatly exceeds the range defined by the parental strains, showing that each parent harbors alleles that are both advantageous and deleterious for this phenotype.

Next, we sequenced the individual recombinant strains and calculated the association between the presence of the C8 variant and virulence at each variant position. To compute P-values, we first computed the t-statistic comparing the group with the C8 allele at that site to the group with the KN99 allele. Rather than comparing this statistic to a theoretical null distribution, we compared it to an empirical null obtained by permuting virulence scores 10,000 times and recalculating the statistics on the permuted virulence levels. For each permutation, we used the most significant t-statistic in the entire genome for the null model; this adjusts for multiple hypothesis testing. Statistics and selected genome-wide P-value thresholds are shown in Figure 3, lower panel. The results from the AA closely mirrored those obtained with BSA, with IR-1 showing the highest statistical significance of any region of the genome. These results validated the BSA approach, which has significant advantages in terms of the number of animals required, the experimental effort, and the potential resolution (see Discussion).

Our results do not imply that all C8 alleles in IR-1 confer a virulence advantage: one or a few alleles in this region could be responsible for the effect, with others being implicated due to linkage (see Discussion). To test the impact of specific alleles, we genetically engineered *C. neoformans* to swap alleles between the two parent strains. To select a pilot locus for this approach, we examined the sequences within IR-1. For protein-coding genes, we looked for the presence of variants which would change one or more amino acids, as an indication of potential perturbation of protein structure or function (Supplemental Table 2). For genes where corresponding deletion mutants were available (39), we assessed these mutants in our mouse model, as an indication of the role that sequence plays in virulence (Supplemental Figure 2, top panel, and Supplemental Table 2). Based on these results, we opted to first test a gene identified as CKF44_03628 that contains two variants, one in the 5’-UTR and another that changes a serine at position 417 to tryptophan. Deletion of this gene resulted in severe hypovirulence (Supplemental Figure 2, top panel, and Supplemental Table 2). To test whether the small nucleotide variants (SNVs) in CKF44_03628 contribute to the virulence difference between C8 and KN99**a**, we engineered reciprocal sequence swaps in the background of each parent strain. Engineering these changes required the introduction of a selectable marker; to control for any marker effect, we transformed each parental strain with both the original sequence and the new sequence (from the other parent) using biolistic transformation with a split marker design (40). We confirmed all strains by whole genome sequencing (WGS; Supplemental Table 1, sheet D) and tested them in mice as above.

Notably, the insertion of a drug marker adjacent to CKF44_03628, with no other sequence change, significantly reduced the lung burden of each parent strain (Figure 4, left, compare KN99 to KN99^**K**^ and C8 to C8^**C**^; see Discussion). Introduction of opposite alleles induced additional changes in lung burden: swapping the C8 allele into KN99 reduced virulence (compare KN99^**K**^ to KN99^**C**^), while the reciprocal change, introducing the KN99 allele into C8, trended towards increased virulence (compare C8^**C**^ to C8^**K**^).

**Figure 4.**
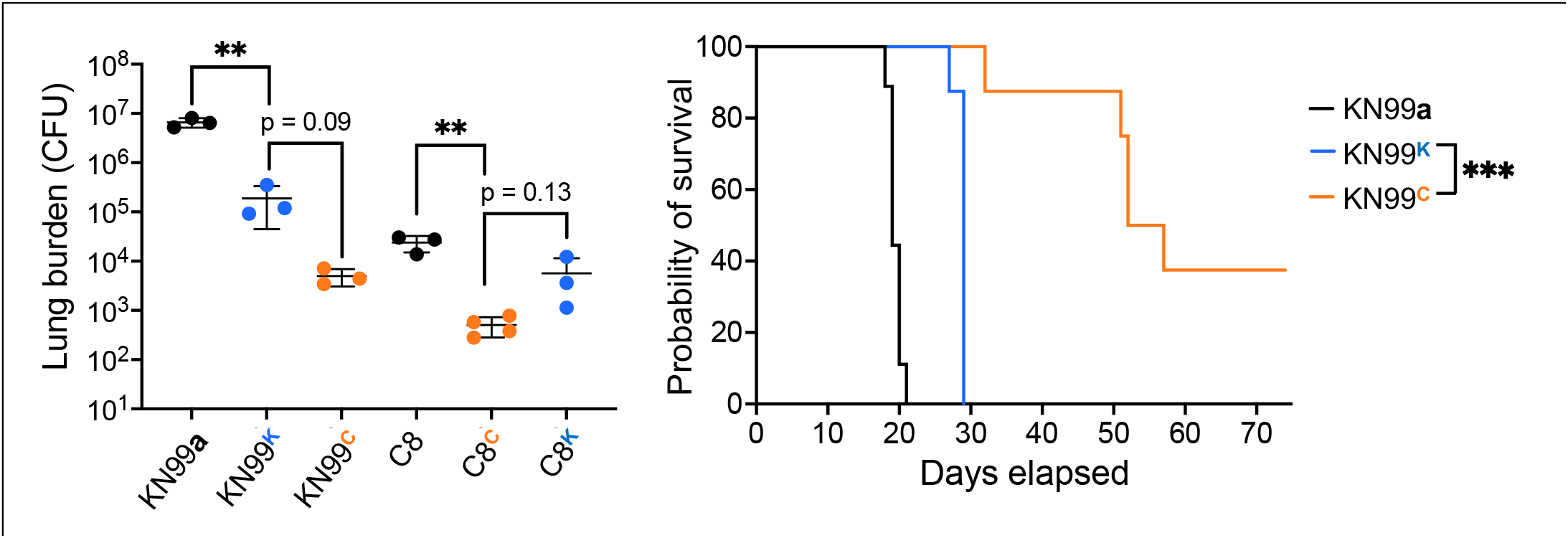
Virulence studies for CKF44_03628 swap strains. Shown are data from intranasal infections of C57BL/6J mice using 12,500 cryptococcal cells of the indicated strain. For this and all subsequent figures, strain names with a superscript have a drug marker inserted near the locus of interest; the base name is the background strain and the superscript indicates the source of the sequence that was swapped into that background (blue, KN99; orange, C8). *Left panel*, total lung burden 9 days after infection with the indicated strain. Each symbol represents an individual mouse; mean and standard deviation are shown. Comparison was by one-way ANOVA with Tukey’s multiple comparisons test *post hoc*. **, p < 0.01. *Right panel*, survival of groups of 8 mice over time after infection, with sacrifice triggered by weight below 80% of peak or signs of disease (see Methods). Survival curve comparison was by Log-rank test with p = 0.0001 for the comparison of KN99^**K**^ to KN99^**C**^.

Consistent with the lung burden results, a survival study with the KN99 background strains (Figure 4, right) showed significantly reduced virulence when the endogenous sequence at this locus was changed to that of C8 (orange), compared to a matched control swap (blue) where the sequence was not altered. All of these results strongly associate the KN99 allele at this position with increased virulence. However, this was the opposite of what we had anticipated for a SNV within IR-1, a region that our BSA and AA studies indicated was associated with increased virulence of the C8 allele. To explain this, we speculated that CKF44_03628, which occurs at the right edge of IR-1, was included in this region because of linkage to one or more C8 alleles with virulence effects large enough to outweigh the KN99 allele advantage at this locus. To test this hypothesis, we looked for a way to refine IR-1.

We had identified IR-1, which is almost 180 kb long and contains 65 genes and 161 variants, by analyzing 100 recombinants from a cross of C8 and KN99. The resolution of such analysis depends on the number of crossovers that occur in the recombinant genomes during meiosis. To refine the boundaries of IR-1, therefore, we needed to increase the number of crossovers in that region. To do this, we attempted to force recombination within IR-1, by performing the same cross with parent strains modified by the insertion of drug resistance cassettes at the ends of IR-1: NAT (41) at the 5’ end in C8 and G418 (42) at the 3’ end in KN99**a**. Selection for progeny able to grow in the presence of both compounds yielded 221 strains (Supplemental Table 1, sheet B), although interestingly, roughly 30% of them showed aneuploidy of chromosome 2 (see Discussion). We used this strain set in a new BSA experiment of the same design as above, although with 87-88 strains per pool, and applied the same criteria we used earlier to identify IR-1 to the results (profiles shown in Supplemental Figure 3). This analysis identified a more limited region of chromosome 2, wholly within IR-1, as implicated in virulence. We termed this region IR-1.1 (Figure 5, top). IR-1.1 spans 57,716 bp (chr2: 297,636 – 355,352) and contains 18 genes and 56 variants (Figure 5, bottom). Notably, it excludes CK44_03628, supporting our conjecture that this locus was implicated in C8 virulence by linkage, rather than because of its inherent properties.

**FIGURE 5.**
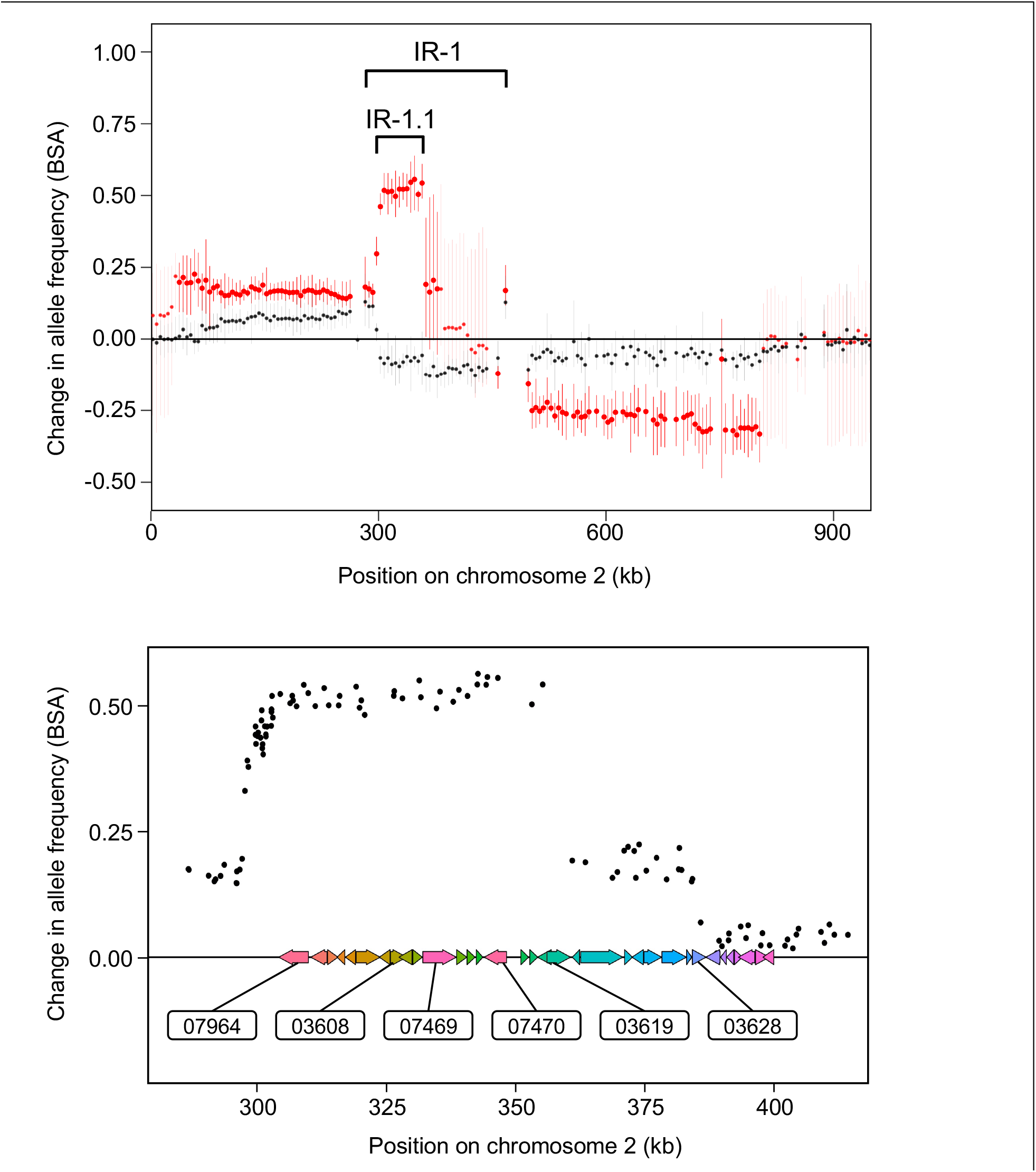
Refinement of IR-1 and genes selected for swap experiments. *Top panel*, BSA analysis of 221 doubly drug-resistant progeny strains from the cross described in the text, performed and presented as in Figure 3. Results for the rest of the genome are in Supplemental Figure 3. *Bottom panel*, BSA results for each individual variant in IR-1 (Supplemental Table 3). Genes are shown as colored arrows; gene identifiers (numerical portion) are shown for the six sequences tested by allele swap experiments.

We next focused our attention on the 18 genes in IR-1.1 (Figure 5, bottom). Based on the effects of variants in this region on protein sequence and the virulence of deletion strains (Supplemental Table 2), we generated reciprocal swap strains for six genes: CKF44_03608, CKF44_03619, CKF44_07469, CKF44_07964, and CKF44_07470 (Supplemental Table 1, sheet D). For genes with more than one variant, we chose transformants where all had been swapped for testing. For the first four of these genes, we observed no differences in virulence between marked parental strains bearing either the original or swapped alleles (Supplemental Figure 4), indicating that none of their SNVs were responsible for the virulence advantage conferred by C8 sequences in IR-1.1. For CKF44_07470, however, we observed increased lung burden at 15 days post-infection in mice infected with marked strains that contained the C8 allele compared to marked strains that contained the KN99 allele (Figure 6, left). Survival studies were consistent with these burden results, showing that KN99 strains with the C8 allele caused more rapid decline of mice than matched strains with KN99 sequence at this locus (Figure 6, right). We saw the same pattern with strains in the C8 background, with higher burden and shorter survival when the C8 allele was present (Supplemental Figures 5 and 6).

**Figure 6.**
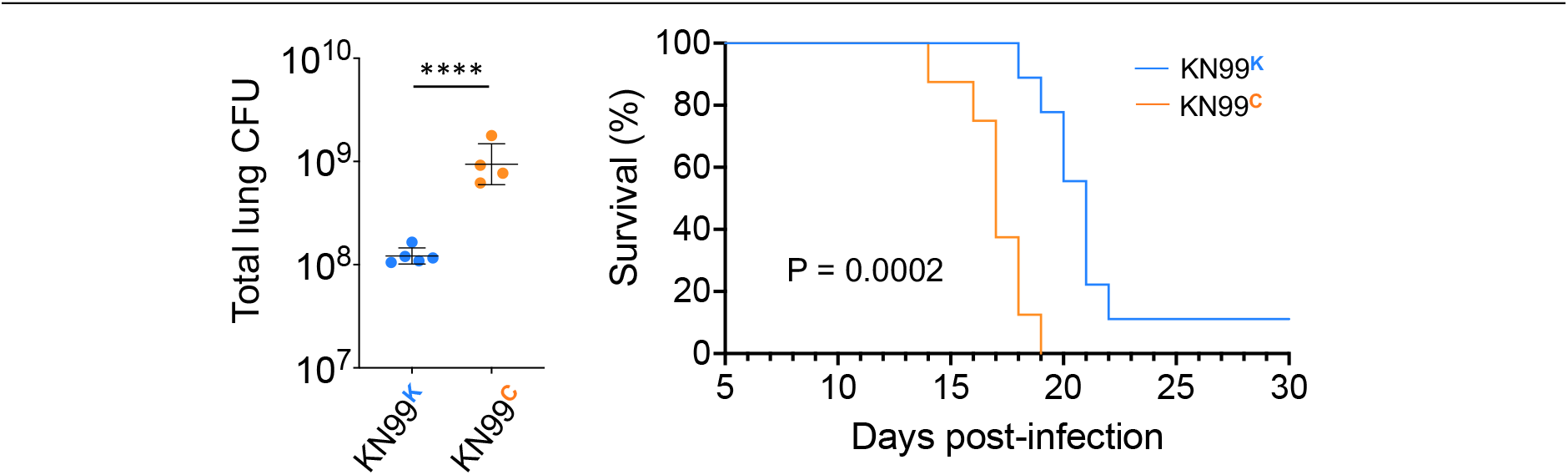
Sequence swaps show that the C8 allele of CKF44_07470 increases the virulence of KN99. Strain designations are as above: the base name indicates the background strain and the superscript indicates the allele that was swapped into that background. *Left*, mean ± SD of lung burden 15 days after mouse infection; each symbol represents one mouse. ****, P ≤ 0.0001 by t-test. Brain burdens are shown in Supplemental Figure 5. *Right*, mouse survival over time, analyzed by Log-rank test.

Our swap studies strongly suggested that one or more C8 variants in the sequence of CKF44_07470 was responsible for conferring a survival advantage on fungal cells in the context of an infected host. This gene, named *PDE2* because it encodes the phosphodiesterase Pde2 (43), contains three differences between KN99 and C8 (Figure 7, top): a 4-nucleotide insertion in C8 in the second intron and two differences in coding regions near the 3’ end of the gene (one synonymous and one missense variant in exon 7). To determine which of these was responsible for the virulence changes we had observed, we separated the variants at each end of the gene by engineering a strain in the KN99 background where the only sequence change in *PDE2* was incorporation of the intron variant (KN99^CKK^). To our surprise, the presence of this variant alone was sufficient to reproduce the increase in organ burden we had previously observed when all variants were swapped (Figure 7, lower left).

**Figure 7.**
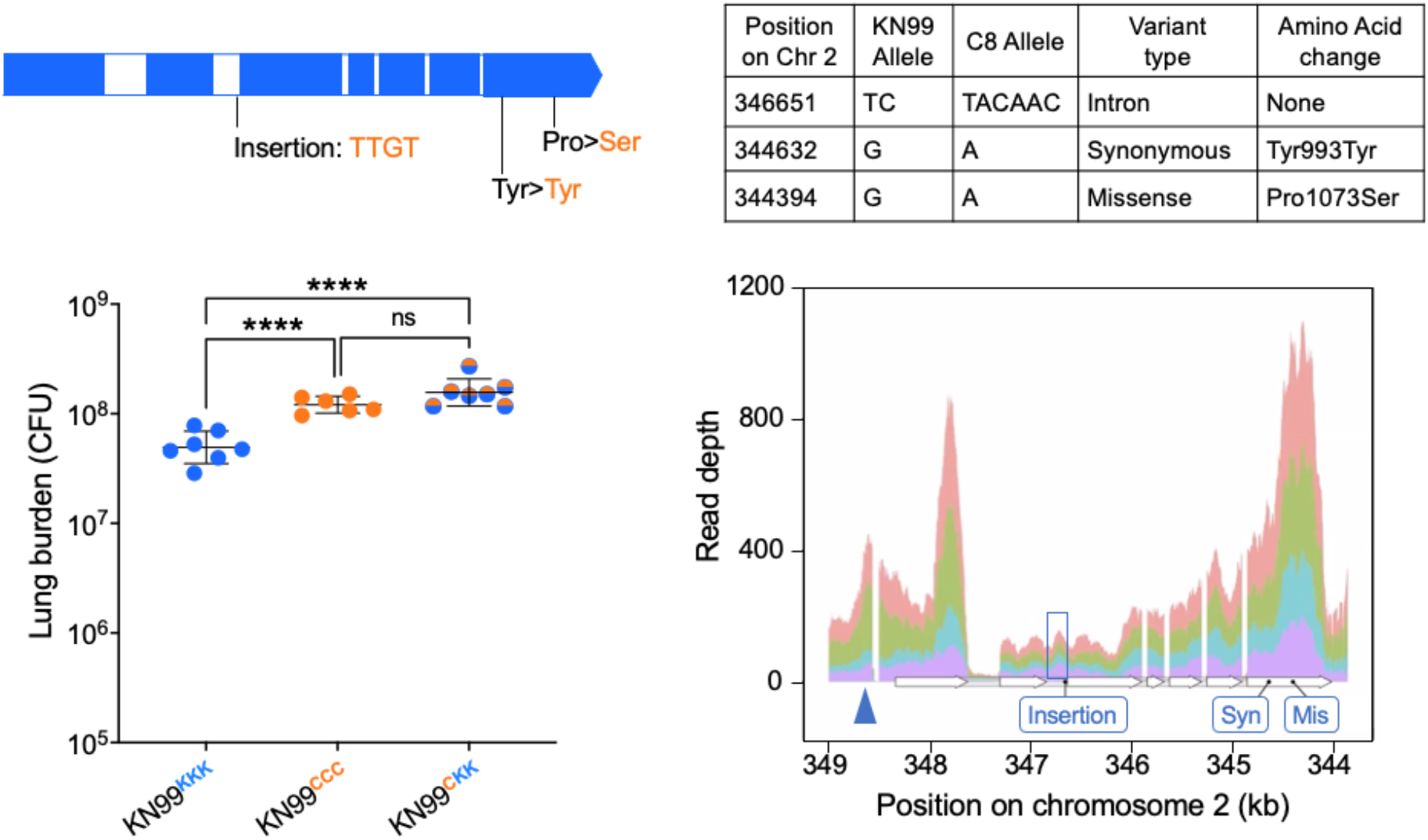
Swapping the intron variant of C8 alone alters virulence as much as all three SNVs. *Top*, variants in C8 relative to KN99. *Lower left*, mean ± SD of total lung burden at 15 days (each symbol represents one mouse). Strain names are as above except that the three letter superscripts represent the three SNVs of CKF44_07470 from 5’ to 3’. Comparison was by unpaired t-test: ns, not significant; ****, p ≤ 0.0001. *Lower right*, read depth from RNA-seq experiments for the chromosome 2 region that contains CKF44_07470 (*PDE2*). Each color represents an independent replicate study using KN99; results for C8 were similar. Below the expression profiles is the current annotation for this region (white arrows, exons; gray segments, intron), with features that differ in C8 versus KN99 sequence indicated in blue. The sequence annotated as the second intron (boxed in blue) is present in mRNA at levels comparable to those of flanking exons, in contrast to all other introns, which show minimal reads. Blue triangle, site of marker insertion.

We wondered whether the 4-bp insertion in intron 2 of C8 might influence virulence by altering the splicing or expression of *PDE2* mRNA. To address this, we performed RNA-seq analysis on each parent strain grown in host-like conditions in vitro (Supplemental Table 1, sheet E). Unexpectedly, we observed robust expression of intron 2 in mRNA from both KN99 and C8, with transcript levels similar to those of the neighboring exons and no indication that it is spliced out during mRNA maturation (Figure 7, bottom right). Based on these results, we concluded that the reference annotation of *PDE2* in H99 and KN99 is incorrect – the region annotated as intron 2 is not in fact an intron. This region in C8 (including the extra 4 bp) is a multiple of three in length, so its presence in the mRNA does not alter the downstream protein reading frame relative to the current annotation. Notably, all 240 non-laboratory sequences that we have analyzed (193 clinical, 4 veterinary, and 43 environmental; see (12) and Supplemental Table 1, sheet A) include this 4-bp sequence. In contrast, the lack of these 4 bp in KN99 shifts the reading frame, resulting in two stop codons (17 and 53 nucleotides downstream of the variant, respectively), so that the encoded protein is truncated 347 amino acids before the predicted active site of Pde2. This likely explains why the lung burden in mice infected with *pde2*Δ is the same as that in mice infected with the parent KN99 (Supplemental Figure 2, lower panel): neither strain produces active Pde2.

Our results suggest that the presence of active Pde2 in C8 has a significant effect on virulence. Since this protein acts to cleave cAMP, we compared the cAMP levels of the two parents. Indeed, cAMP was significantly higher in KN99 than in C8, consistent with the lack of an active phosphodiesterase (11.8 ± 1.05 versus 4.19 ± 0.33 pg/10^6^ cells; P <0.0001 by t-test).

## DISCUSSION

We set out to discover, in an unbiased way, naturally occurring sequence variants in the *C. neoformans* genome that influence virulence. Our goal is to identify new sequences of interest, which can become a focus of direct research attention. By working at nucleotide resolution, we can gain mechanistic understanding of how virulence is influenced by specific sequence changes, whether they occur in novel or previously characterized genes. Our strategy further has the potential to identify key variants located in regulatory elements or essential genes.

We used two approaches to discover sequence variants of interest, applying both to a population of recombinant progeny derived from a cross between a clinical strain (C8) and a laboratory strain (KN99). One approach was bulked segregant analysis (BSA), for which we compared genome sequence of recombinant pools either grown non-selectively *in vitro* or recovered from mouse lungs after intranasal inoculation. The second was to analyze the association between sequence and virulence, with the latter measured by fungal lung burden in the same animal model. The two methods showed excellent agreement in identifying a region of chromosome 2 where the C8 sequence favored higher lung burden compared to the same region of KN99. We then used sequence swap experiments to identify a gene within this region that increased the virulence of C8, and, ultimately, to narrow the key region to a single 4nucleotide difference.

The power of our initial analysis came from combining two completely distinct methods. Association analysis was based on infecting mice with individual *C. neoformans* strains, with a readout of lung burden determined by plating lung homogenates. In contrast, BSA was based on infection with pooled strains, with a readout of allele frequency determined by whole genome sequencing. BSA has considerable methodological advantages. First, it uses fewer mice (up to two orders of magnitude), which is of both ethical and practical significance. Second, due to the efficiency of pooling strains and the capacity of genome sequencing, BSA requires less experimental work, even beyond that related to the animal studies. For these reasons, once we established the robust agreement between the two methods, we moved to BSA analysis alone.

Drug resistance markers are a convenient tool for strain engineering in *C. neoformans* (41,42,44). However, insertion adjacent to genes of interest may reduce baseline strain virulence (45), as occurred when we inserted a marker upstream of CKF44_03628 (Figure 4). We hypothesize that this is because of interference with unrecognized regulatory sequences, a conjecture supported by our RNA-seq data (Supplemental Table 1, sheet E). We also used marker cassettes at each end of IR-1 in the parental strains to force recombination within this region. While this strategy was successful, our analysis suggests two cautionary notes. First, in addition to enhanced recombination between marker sites, these studies yielded a high level of aneuploidy of this chromosome, likely due to the pressure imposed by double drug selection. In principle, the frequency of these strains in the starting or final pools should not affect the difference in frequency between alleles, since each aneuploid strain contains both alleles; aneuploidy may thus reduce statistical power but should not lead to P-value inflation. Our analysis of the same strain set with and without the aneuploid progeny indeed confirmed that their presence did not alter our results. However, in applications where the presence of aneuploids is a concern, strains generated by ‘forced’ recombination in this manner should be subjected to whole genome sequencing and analysis of copy number variation. Second, a recombination hotspot between our two sites of marker insertion yielded multiple recombinants with the same breakpoint. For future studies, it would be advantageous to avoid such insertion sites, to obtain more evenly distributed recombination events.

Our studies of *C. neoformans* recombinants show that each parental strain harbors multiple significant variants that influence virulence in both directions. For crosses of C8 and KN99, the genomic region with the largest change in allele frequency during infection happened to be one in which the C8 sequence favored higher lung burden, even though the C8 strain overall is less virulent than KN99. When we investigated individual sequences within this region, we found genes where the C8 allele favored virulence (e.g. *PDE2*), the KN99 allele favored virulence (e.g. CKF44_03628), or neither favored virulence (e.g. CKF44_03619). All these patterns can occur within the same identified region because the sequences are genetically linked. Our refinement of IR-1 yielded more concordant patterns, with the exclusion of CKF44_03628 from the region where C8 favored virulence. In theory, using enough recombinants would allow resolution of individual genes; in practice, experimental factors will dictate the balance between the effort expended to generate and analyze recombinants and the effort required to dissect an identified region of interest through allele swapping. Such factors may include the mating efficiency of specific strains and the efficiency of cryptococcal genome engineering, which has recently advanced through use of CRISPR (46,47).

The gene we identified in IR-1 for which the KN99 allele favored virulence, CKF44_03628, is robustly expressed in human CSF (48,49) and in multiple *in vitro* conditions relevant to virulence (50). This gene encodes a homolog of the *S. cerevisiae* Vps45, which acts in the regulation of vesicular transport (51). Cells completely lacking Vps45 were recently characterized in *C. neoformans* and shown to be impaired in iron uptake, mitochondrial function, and surface properties that are key factors in virulence (52). Our virulence studies suggest that the C8 variants in this sequence compromise Vps45 activity; future examination of these swap strains could potentially define this mechanistically.

We were surprised that our virulence-based analysis yielded *PDE2*, since this gene had been reported to play a minimal role in cryptococcal expression of virulence factors (43) and a deletion strain from the Madhani collection (39) showed no altered virulence in our animal model. This mystery was solved by our discovery that the *PDE2* transcript in the laboratory strains used for these studies does not encode an active protein, so deletion of the gene would not alter phenotype. Consistent with this finding, the level of cAMP in KN99 is significantly higher than that of the clinical strain C8. Nonetheless, we cannot rule out the possibility that Pde2 plays additional cellular roles, independent of cAMP.

Although the cryptococcal literature suggests that higher cAMP generally favors the development of virulence factors, cAMP levels are subject to complex regulation through multiple pathways (53,54) and compensatory changes in KN99 may mitigate the effects of Pde2’s absence. Feedback mechanisms may also participate, as suggested by increased transcription of *PDE2* in KN99 compared to C8 in hostlike conditions (Supplemental Table 1, sheet E), even though this mRNA does not encode a functional phosphodiesterase. Overall, the common association of higher cAMP levels with virulence may need to be refined.

We suspect that the defective KN99 allele appeared near the time of isolation of its progenitor strain H99 (33), because it occurs in all lineages originating in this isolate (8,26) but not in any of the 240 clinical or environmental isolate sequences that we examined. This is the second virulence-altering genomic change that has now been associated with common laboratory strains of *C. neoformans*; prior studies showed that one branch of the H99 lineage, which includes the most prevalent model strains, has increased virulence due to partial deletion of *SGF29* (26,55). These differences must be kept in mind as future genome-wide projects are pursued in this organism; while well-developed reference strains and their derivatives are a tremendous tool for research, they do not always accurately represent clinical isolates (56).

Our strategy, and the libraries of recombinants we have collected, are powerful tools for the unbiased analysis of cryptococcal traits of biological and medical interest at the sequence level. Any characteristic of interest that is measurable and varies within the recombinant population is amenable to this analysis, even if it is fairly similar in the original parents. Such studies, in our lab and others, will identify new targets for investigation and lead to increased mechanistic understanding of an important fungal pathogen of humans.

## ACKNOWLEDGEMENTS

We thank John Perfect, Andrej Spec, and the CINCH consortium for generously providing clinical strains. We appreciate the assistance of Chase Mateusiak with computational analysis, Alyssa Brunsmann with mouse studies, Abigail Kimball with computational method assessment, and Elizabeth Nordmark, Gaby Altman, and Michael Lin with spore isolation. We thank Barak Cohen for suggesting forced recombination, members of the Doering lab for stimulating discussion, and Keeley Choy, Ellie Gaylord, Daphne Ko, and Liza Loza for comments on the manuscript.

## AUTHOR CONTRIBUTIONS

D.P.A.: Conceptualization, Methodology, Software, Validation, Formal Analysis, Investigation, Data Curation, Writing – Original Draft, Writing – Review & Editing, Visualization. H.L.B.: Methodology, Investigation. G.C.: Methodology, Investigation. M.R.B. Conceptualization, Methodology, Writing – Review & Editing, Supervision, Funding Acquisition. T.L.D: Conceptualization, Writing – Original Draft, Writing – Review & Editing, Visualization, Supervision, Project Administration, Funding Acquisition.

## METHODS

### Strains and growth

KN99 strains were obtained from Joe Heitman (Duke University) and clinical strains from John Perfect (Duke University) and from the CINCH consortium; see Supplemental Table 1 for details. *C. neoformans* deletion strains generated by the Madhani group (39) were obtained from the ATCC. For experiments, strains were streaked from storage at -80 °C onto YPD plates, incubated for two days at 30 °C, inoculated from single colonies into YPD liquid cultures, and grown at 30 °C with shaking (220 rpm) unless indicated otherwise.

### Virulence studies

For all infections, overnight cultures of cells grown as above were washed three times in PBS, adjusted to 2.5 × 10^5^ cells/ml, and used for intranasal inoculation of 6-week-old female C57/Bl6 mice (see Figure legends for specific inocula) and for plating to confirm viable cells in the inoculum. To measure organ burden, mice were sacrificed at the times indicated in the text and organs were harvested, homogenized, and plated (YPD agar, 30 °C, 2 days) for enumeration of colony forming units. To assess survival after infection, mice were weighed daily and sacrificed if their weight reached 80% of initial weight, if they showed signs of illness, or at the end of the study. Statistical differences in organ burden and survival were assessed by one-way ANOVA with Tukey’s multiple comparisons test *post hoc* and Logrank (Mantel-Cox) test, respectively, using GraphPad Prism9.

### Crosses and spore isolation

To assess filamentation, single colonies of strains to be tested were mixed with single colonies of either KN99**a** or KN99α on V8 medium (per liter: 50 ml V8 juice, 25.7 ml 0.2M Na_2_HPO_4_, 24.3 ml 0.1 M citric acid, 40 g agar, and 2 ml 25 mM CuSO_4_ (added after autoclaving)) and incubated for 14 days before examination on a dissecting microscope. For individual strains that filamented poorly under these conditions, additional test crosses were performed on V8 plates adjusted to pH 7.0 by using 0.5 g KH_2_PO_4_ in place of the Na_2_HPO_4_ and citric acid solutions. For drug selection of recombinant progeny, cells were similarly crossed, and progeny plated on double drug plates. Colonies were then passaged three times on drug plates, once on YPD, and frozen in YPD. For spore microdissection, cells of each parent were grown as above, washed twice in PBS, diluted to OD_600_ of 1, and spotted on V8 plates (medium made as above but passed over a 70 μm pore strainer before plating).

### Genome sequencing and analysis

For gDNA isolation, cells were either grown overnight in YPD as above or recovered from BSA studies as below. DNA was then isolated and sequenced as in (32), except that sonication was to an average size of 300 bp and sequencing was on an Illumina HiSeq-2500 (for paired end 150-bp reads). Reads were aligned to the KN99α ASM221672 reference genome (32) using NextGenMap (57) with the -X 100000000 parameter. Output SAM files were converted to BAM and PCR duplicates were removed using the `view` and `rmdup` commands from SAMtools 1.7 (58), respectively. Unmapped and soft clipped reads (with at least 20 nucleotides) were extracted using the split_unmapped_to_fasta.pl script from the Lumpy package (59) with the -b 20 parameter and realigned using the split-read aligner Yaha (60) with the -M 15 -H 2000 -L 11 parameters. The outputs of NextGenMap and Yaha were merged using the script MergeSamFiles from Picard tools version 2.10.0 (https://github.com/broadinstitute/picard) and indexed with the index function from SAMtools.

### Variant calling

SNVs and small indels were identified using FreeBayes version 1.1.0 (61) with the parameters -F 0.75 -! 5 -p 1 -m 30. Variant annotation of the resulting VCF files was performed with SNPEff version 4.3.1 (62). Copy-number variants (CNVs) were called with CNVnator 0.3.2 (63) using default parameters. As in (32), each CNV was assigned a mean depth of coverage relative to the genome-wide depth of coverage and considered to be a duplication if the normalized depth was at least 1.9 times the genomewide average for the same strain or a deletion if the normalized depth was below 0.25 times the genome-wide average.

### Bulked segregant analysis (BSA)

For BSA, strains for analysis were streaked from frozen stocks as above, inoculated into 600 μl of YPD in 96-well deep-well plates, covered with Breathe-Easy film, and grown overnight (30 °C, 500 rpm). Equal volumes from each well were combined as specified below, in biological duplicate, and aliquots of the resulting pools were used for three purposes as follows: (1) immediately reserved to represent the initial pool; (2) grown in YPD (5 × 10^4^ cells in 4 ml, 30 °C, 220 rpm, 20 h); (3) used to infect mice as above. After 9 or 15 days, lungs were harvested and homogenized in 3 ml of DNase I buffer (10 mM Tris-HCl, 2.5 mM MgCl_2_, 0.g mM CaCl_2_, pH 7.5) containing 2 mg/ml DNAse I (Thermo Scientific) and the homogenate was filtered to remove host tissue fragments using a cell strainer with 40 μm pores. The filtrate was subjected to centrifugation (3000xg, 7 min, RT) and the pellet was resuspended and incubated for 5 minutes in 2 ml of 1% SDS to lyse remaining host cells. The suspension was then diluted 10-fold in distilled water, centrifugation and SDS incubation were repeated, and the fungal cells were washed twice with distilled water.

DNA was prepared from all samples and sequenced using Illumina technology as above (average coverage 100.2-fold). Reads from replicate pairs were combined and the number of reads matching each parent’s allele at each variable site was used as an estimate of allele frequency. We next calculated the change in allele frequency at each variable site for growth in YPD or in mice relative to the initial pool sample. We also calculated a genome-wide significance P-value with the R package QTLseqr (64), applying the G’ method (38) to each pool individually and to the combined reads from all pools. Data was plotted as changes in allele frequency, with smoothing as in reference (64) when indicated in the text. BSA #1 included 100 strains derived from a KN99**a** × C8 cross, randomly assorted into 5 pools of 40 strains each, with each strain present in 2 pools and each pair of pools sharing 10 strains. BSA #2 included 221 new strains derived from a KN99**a**-G418 × C8-NAT cross. These strains were divided in 5 pools containing 87-88 strains each, with each strain present in 2 different pools. A sixth pool included all of the strains together.

### Association Analysis (AA)

Individual mice were infected with KN99 and sequenced recombinant strains and CFU were assessed at 9 days as above, using lung burden (log_2_ fold-change in CFU compared to KN99) as a surrogate for virulence. At each variant position in the genome, we compared the virulence of strains with either KN99 or C8 alleles by calculating the t-statistic. Null distributions for P-values were obtained by permuting the assignment of CFU phenotypes to strains 10,000 times, calculating the largest t value genome-wide, and using the distribution of those largest t-values. This method accounts for multiple hypothesis testing.

### Strain engineering

For sequence swap experiments, we engineered strains using biolistic transformation and the split marker strategy described in (40). Selection was mediated by a nourseothricin resistance marker (41) that was inserted adjacent to the gene of interest (GOI) at the end nearest the variants to be altered, avoiding putative promotor or terminator regions. One fragment used for transformation therefore included one end of the marker gene fused to flanking and coding region of the GOI by PCR; the other consisted of the rest of the marker gene (including a 271 bp overlap) and sequence farther away from the GOI. Details of specific strain constructions are available on request. All candidate transformants were assessed by WGS as above to select strains that had undergone recombination to yield the desired change in variant.

### RNA seq and analysis

RNA-seq was performed as in Reuwsaat *et al* (65) with minor differences. RNA was isolated from parental or engineered strains grown for 24 hours in RPMI + 10% mouse serum (37°C, 5% CO_2_) and sequenced as previously described (66). Briefly, cDNA samples were sequenced using the Illumina Nextseq platform for paired-end 2 × 150 bp reads and read quality was evaluated by FastQC (67). Fastq files were aligned to the KN99α genome (32) using Hisat2 version 2.2.1 (68) with the default parameters plus --max-intronlen 2200. SAM files were converted to bam, reads were sorted and indexed, and read duplicates were removed from the final bam files using SAMtools 1.7 (58). The number of reads mapped per gene was calculated using featureCounts from the package Subread 2.0.0 (69) and differential gene expression was analyzed with DESeq2 (70), using the IHW (independent hypothesis weighting) package to calculate adjusted p-values (71).

### cAMP analysis

Strains were grown as above, washed in RPMI with 2% mouse serum that had been preconditioned at 37 °C and 5% CO_2_, resuspended in 20 ml of the same medium at 10^7^ cells/ml, and grown in the same conditions for 24 h. Cells were then again counted, and 1-ml aliquots were collected by sedimentation, flash frozen, and submitted to the Washington University School of Medicine Metabolomics Facility for liquid chromatography/mass spectrometry analysis of cAMP relative to a ^13^C-5-adenosine cAMP internal standard (Toronto Research Chemicals).

## SUPPLEMENTAL TABLES

**Supplemental Table 1**. Strain and sequence information. *Sheet A*, clinical isolates; *Sheet B*, parental and recombinant strains; *Sheet C*, BSA data; *Sheet D*, engineered swap strains; *Sheet E*, RNA-seq data.

**Supplemental Table 2**. Details of genes within IR-1, including virulence phenotypes.

**Supplemental Table 3**. All variants within IR-1.

## SUPPLEMENTAL FIGURES

**Supplemental Figure 1**. Unmarked recombinant BSA results for all chromosomes.

**Supplemental Figure 2**. Virulence studies of deletion strains.

**Supplemental Figure 3**. Marked recombinant BSA results for all chromosomes.

**Supplemental Figure 4**. Lung burden data for sequence swaps that did not influence virulence.

**Supplemental Figure 5**. Brain burden data for CKF44_07470 swaps in C8 and KN99 background.

**Supplemental Figure 6**. Lung burden and survival for CKF44_07470 swaps in C8 background.

