## Supplemental Figures 1-6 for "Unbiased discovery of natural sequence variants that influence fungal virulence"

**SUPPLEMENTARY FIGURE 1**

BSA analysis of 100 unmarked C8 x KN99 progeny strains. Shown are changes in allele frequency for experimental samples compared to initial pools, with positive and negative values arbitrarily assigned to changes in the direction of C8 and KN99, respectively. Values were plotted for 5-kb windows as in Figure 3, but without smoothing and with a false discovery rate cutoff of 0.05. Symbols, mean value for all pools; vertical lines, range of individual pool values; black, YPD samples; red, mouse lung samples.

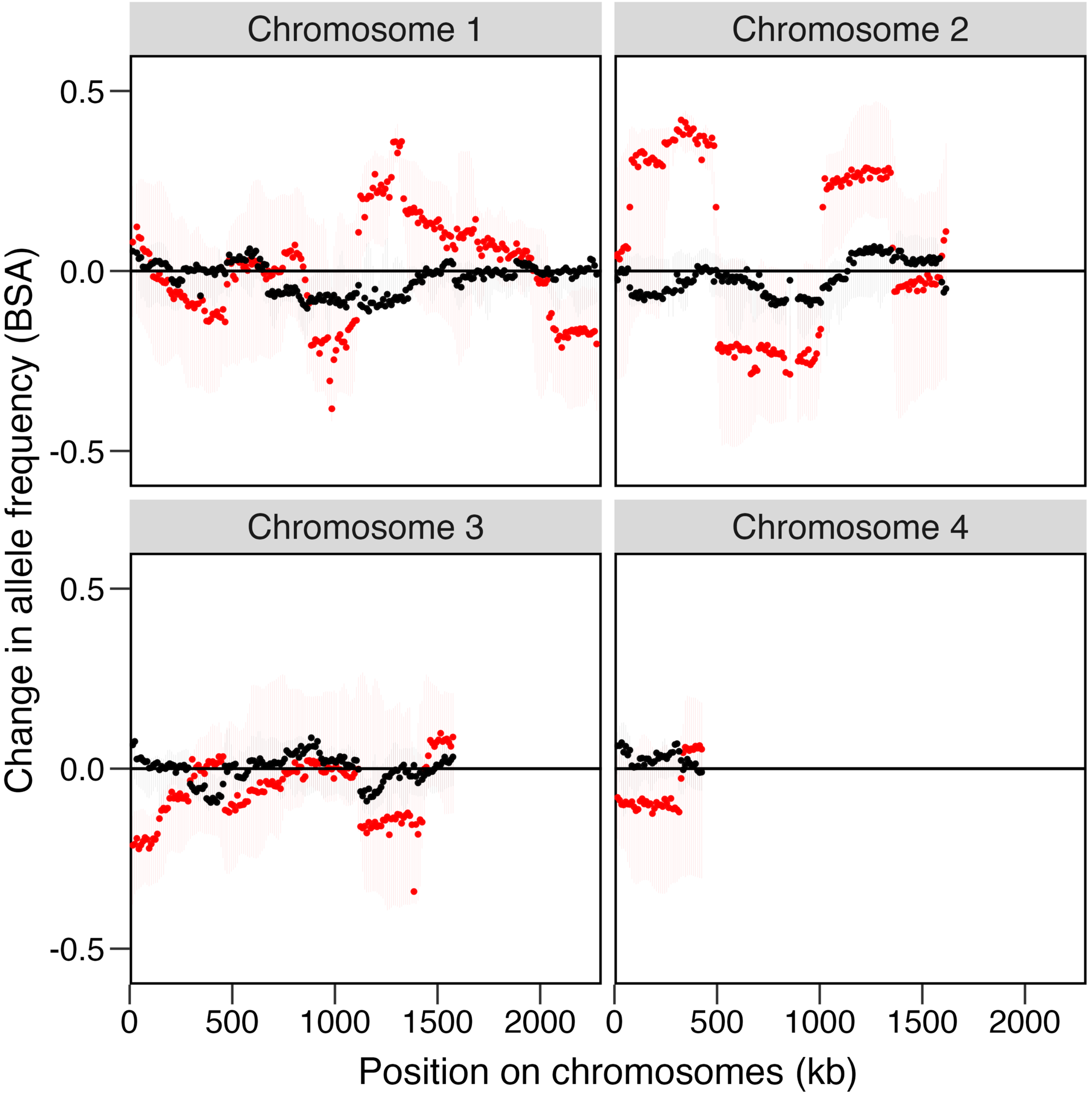

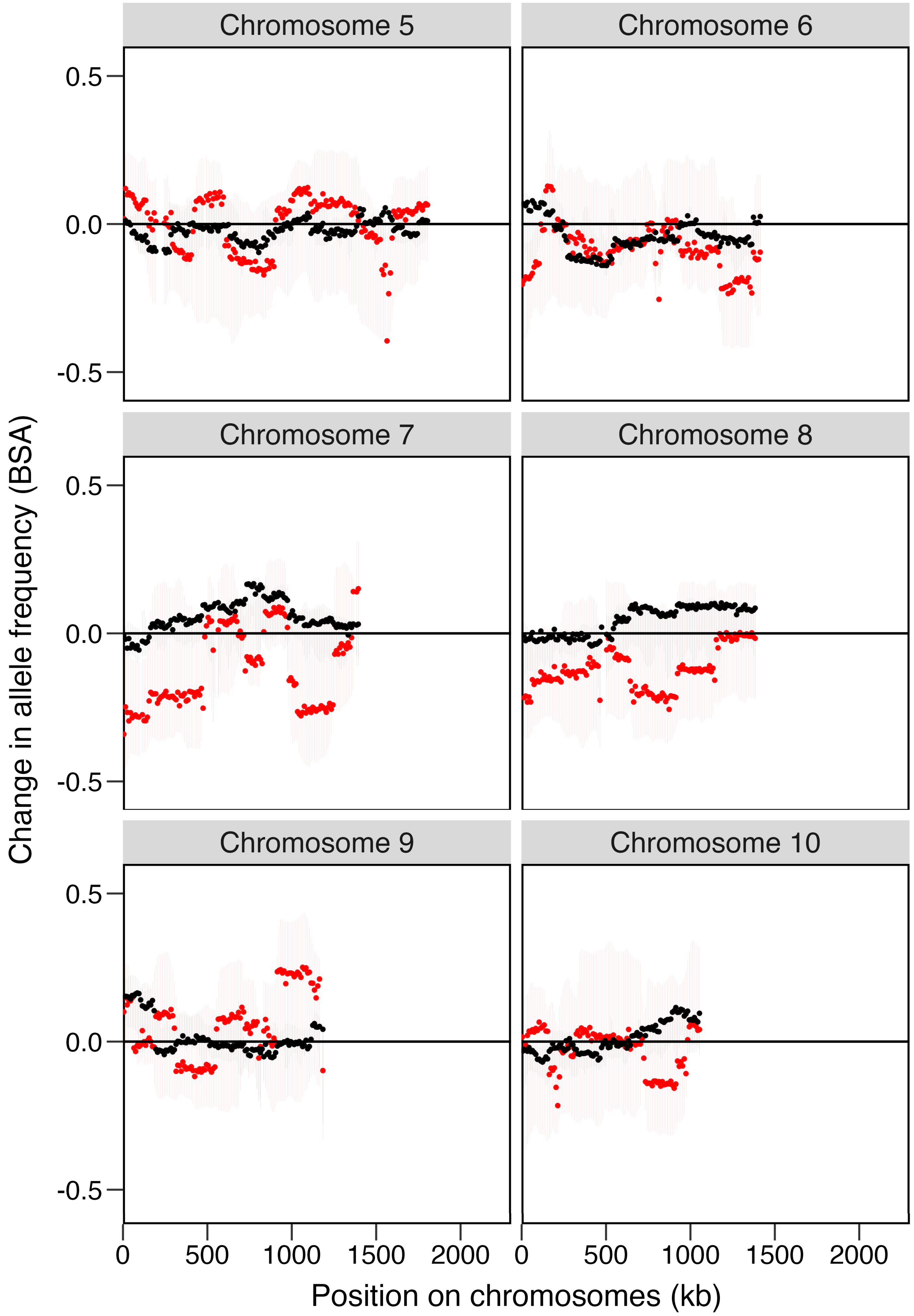
Chromosome 5
Chromosome 6
Chromosome 7
Chromosome 8
Chromosome 9
Chromosome 10
0.5
0.0
-0.5
Change in allele frequency (BSA)
0
500
1000
1500
2000
Position on chromosomes (kb)

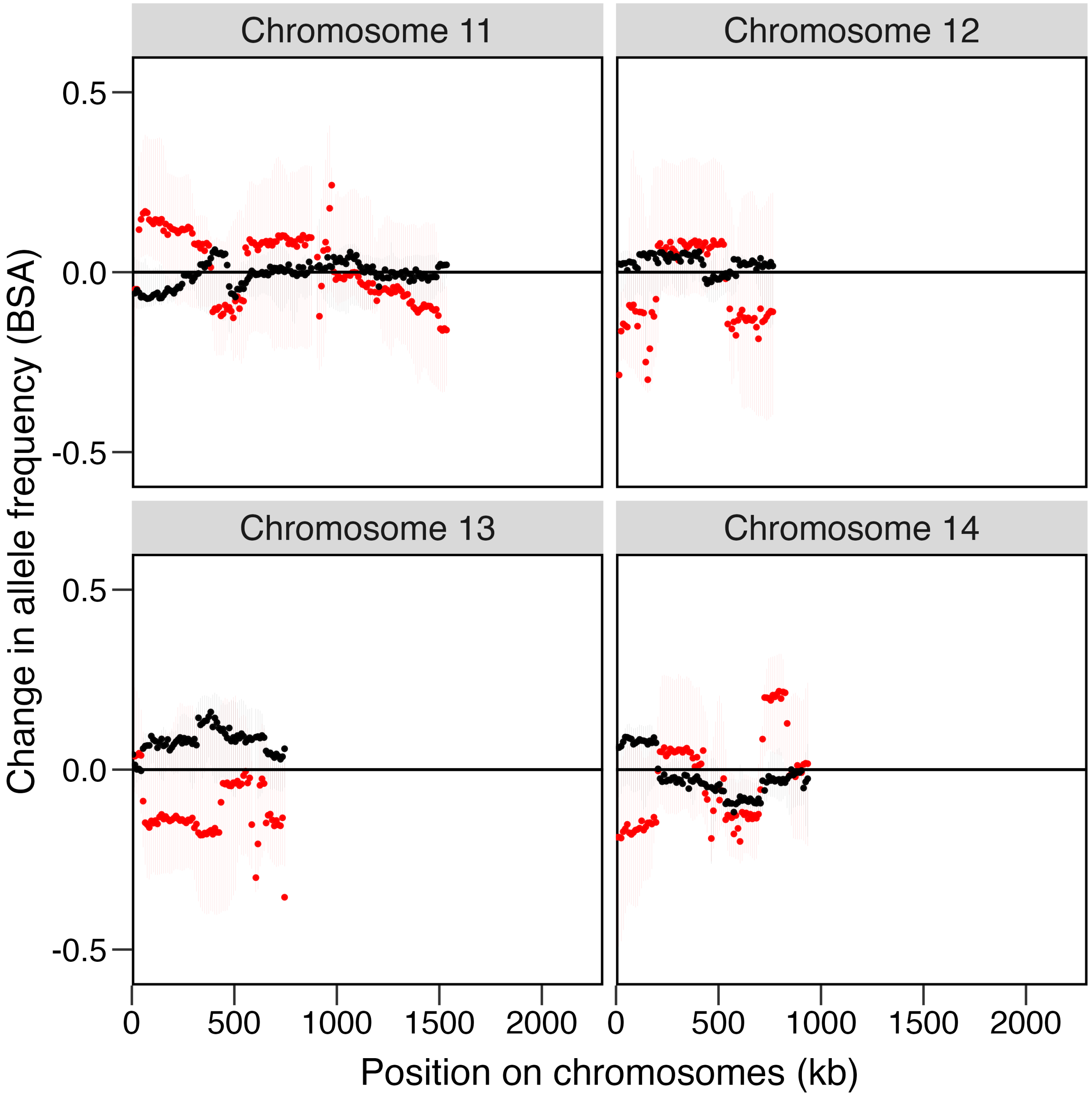

Chromosome 11
Chromosome 12
Chromosome 13
Chromosome 14
0.5
0.0
-0.5
0
500
1000
1500
2000
Change in allele frequency (BSA)
Position on chromosomes (kb)

**SUPPLEMENTARY FIGURE 2**

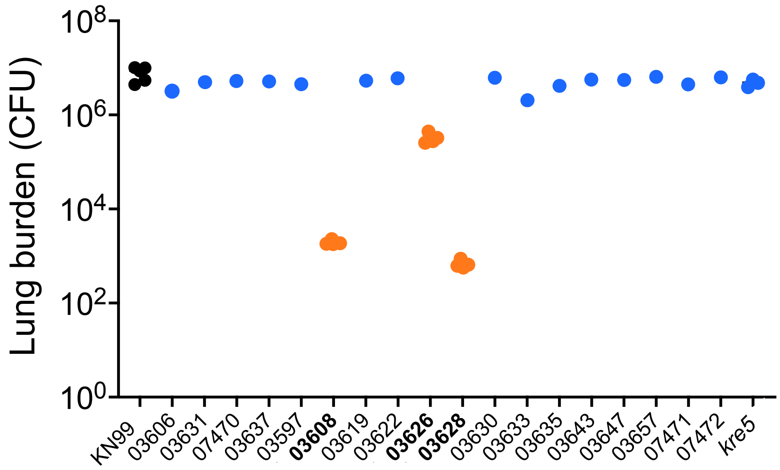


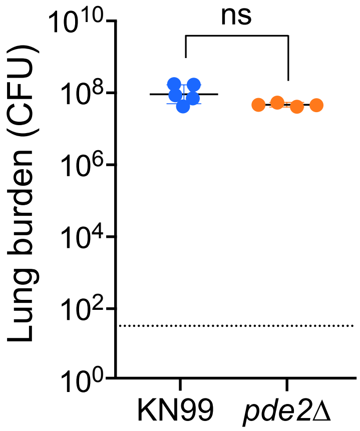


**Supplementary Figure 2.** Virulence of mutant strains, as assessed by total lung burden. C57Bl/6J mice were intranasally inoculated with 12,500 cells of KN99 or strains in the same background lacking the indicated genes (denoted by name or the numerical portion of the CKF44 identifier). Each symbol corresponds to an individual mouse. *Top panel*, mutants lacking genes in IR-1, assessed at 9 days post-infection. After an initial study comparing one mouse per mutant to a group of five infected with KN99, strains that differed from WT (indicated in bold) were tested in three additional animals; the combined results are shown. *Bottom panel*, lung burdens of *pde2*Δ and KN99 at 15 days post-infection. Dotted line, limit of detection.

### SUPPLEMENTARY FIGURE 3

BSA analysis of 221 doubly drug-resistant progeny strains from the cross described in the text. Shown are changes in allele frequency for experimental samples compared to initial pools, with positive and negative values arbitrarily assigned to changes in the direction of C8 and KN99, respectively. Values were plotted for 5-kb windows as in Figure 3, but without smoothing. Symbols, mean value for all pools; vertical lines, range of individual pool values; black, YPD samples; red, mouse lung samples. Larger symbols and darker lines indicate regions of statistical significance, as defined in the text but with a false discovery rate cutoff of 0.05.

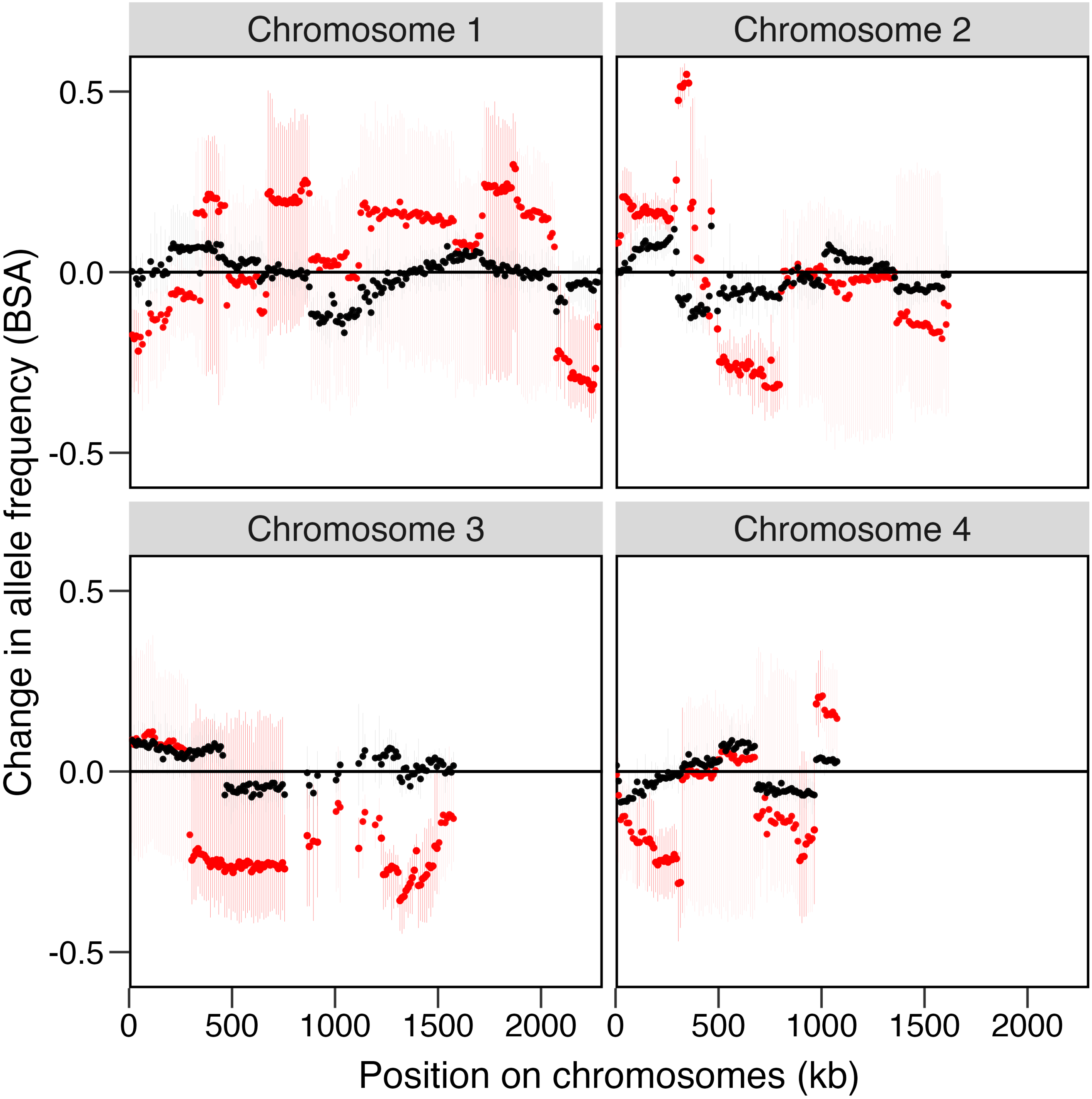

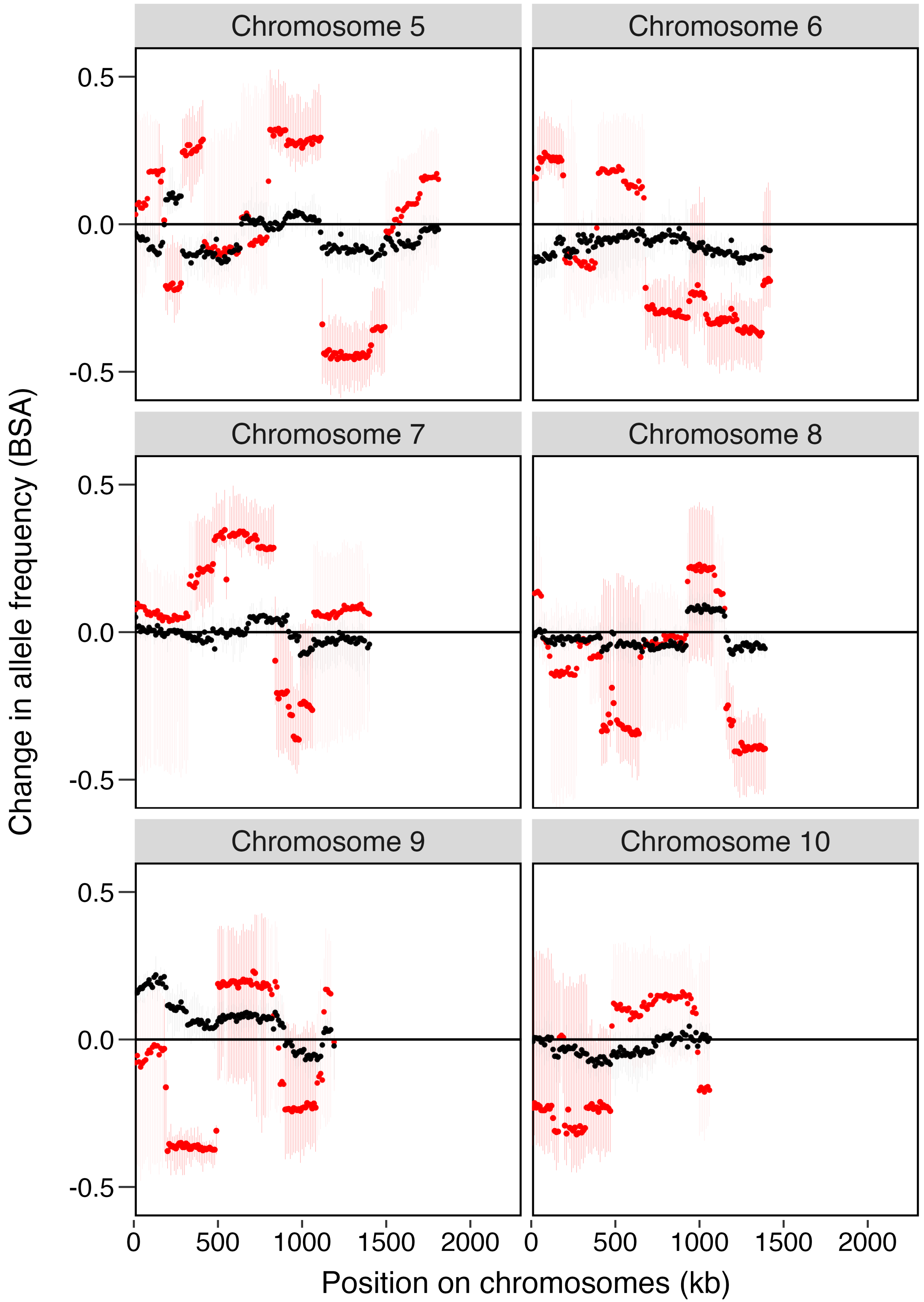

Chromosome 5
Chromosome 6
Chromosome 7
Chromosome 8
Chromosome 9
Chromosome 10
0.5
0.0
-0.5
0
500
1000
1500
2000
Change in allele frequency (BSA)
Position on chromosomes (kb)

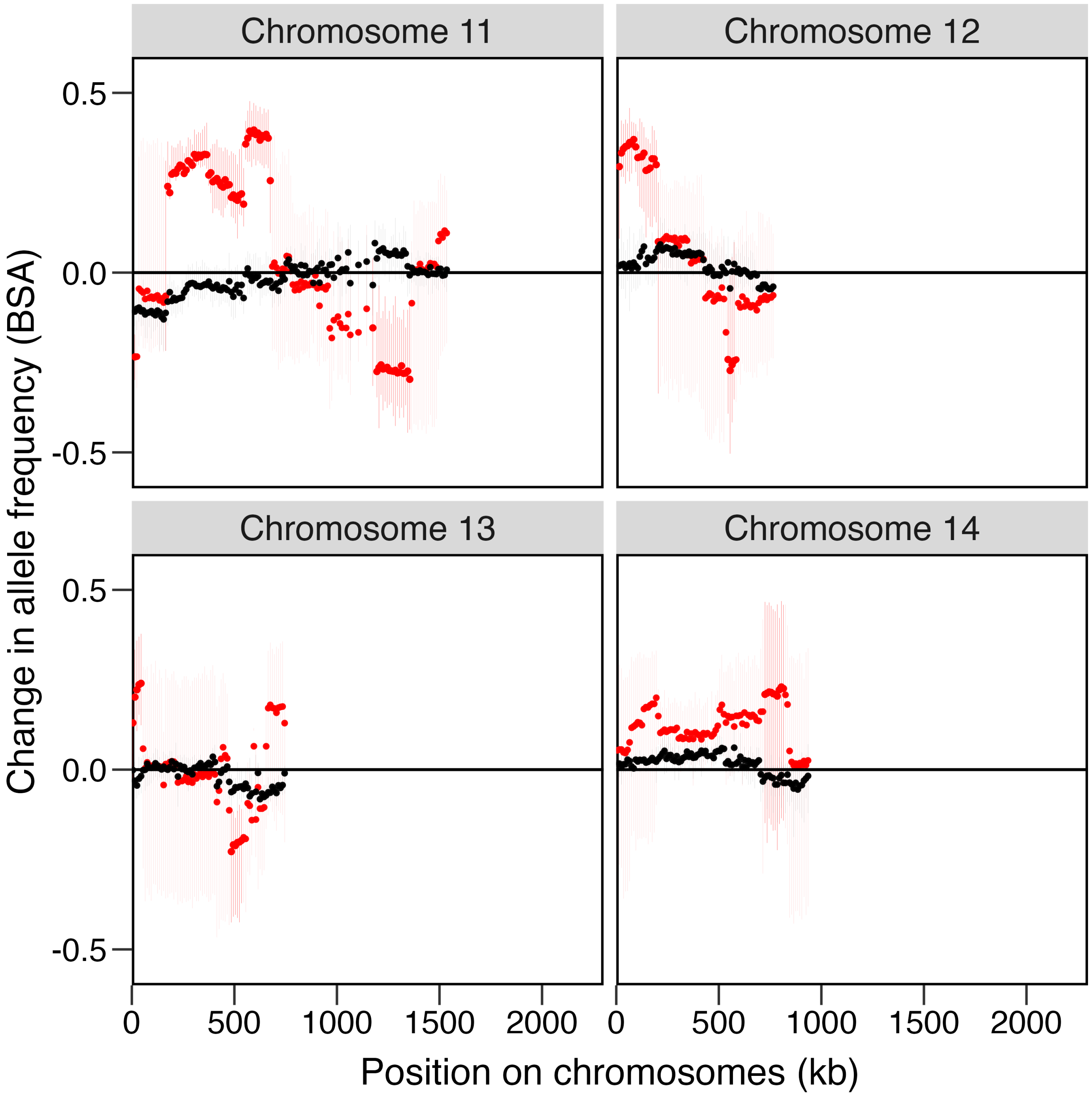
Chromosome 11
Chromosome 12
Chromosome 13
Chromosome 14
0.5
0.0
-0.5
0
500
1000
1500
2000
Change in allele frequency (BSA)
Position on chromosomes (kb)

### SUPPLEMENTARY FIGURE 4

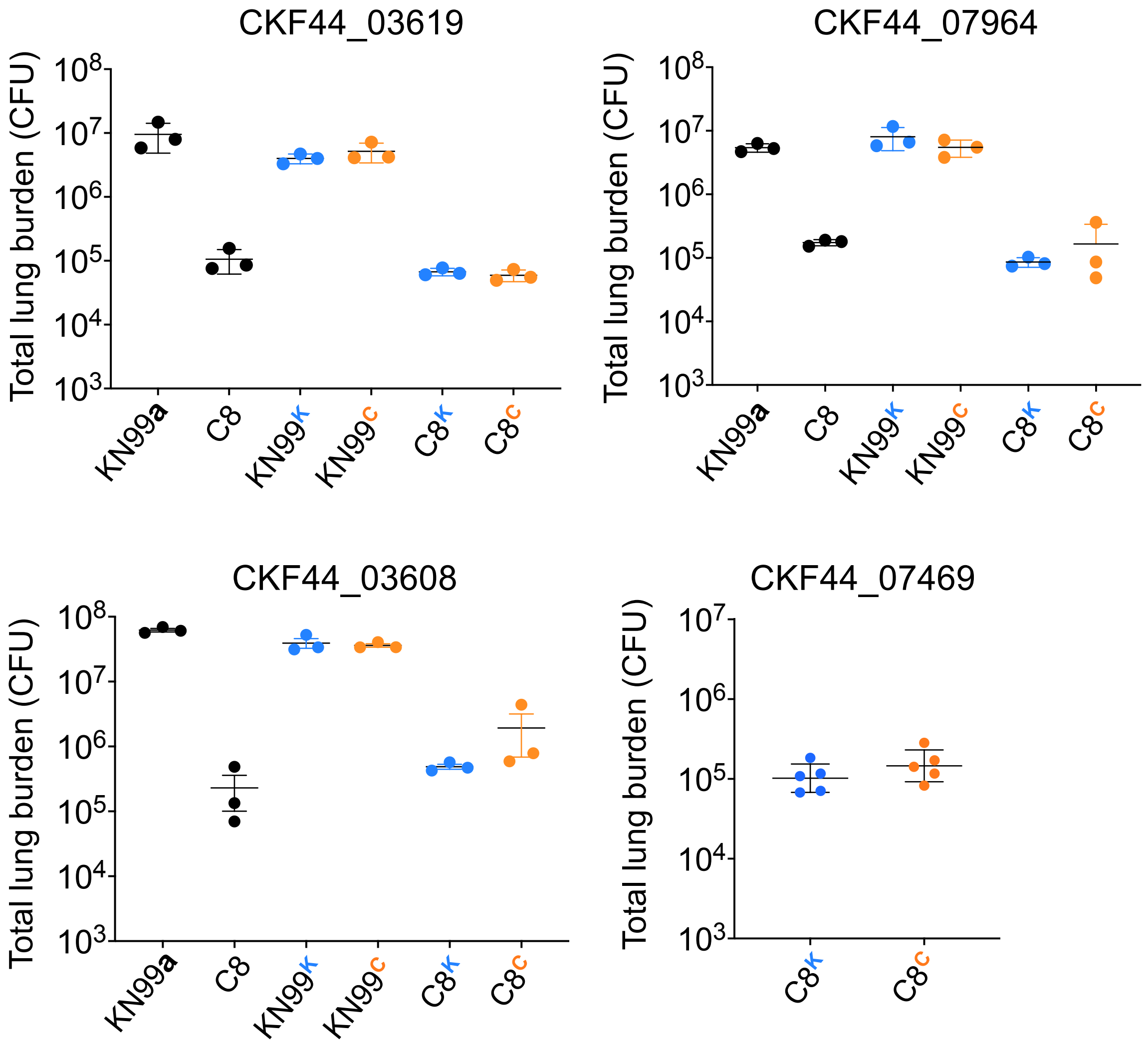


**Supplementary Figure 4.** Allele swaps between C8 and KN99 of the four genes indicated above do not alter virulence. Strain designations are as in the text: the base name indicates the background strain and the superscript, if present, indicates the allele that was swapped into that background. Details of the swaps are in Supplementary Table A, sheet 4. Infection and measurement of lung burden were as in the Methods and mean ± SD of total lung burden is shown; each symbol represents one mouse. None of these experiments showed any significant differences in lung burden between strains in the same background.

**SUPPLEMENTARY FIGURE 5**

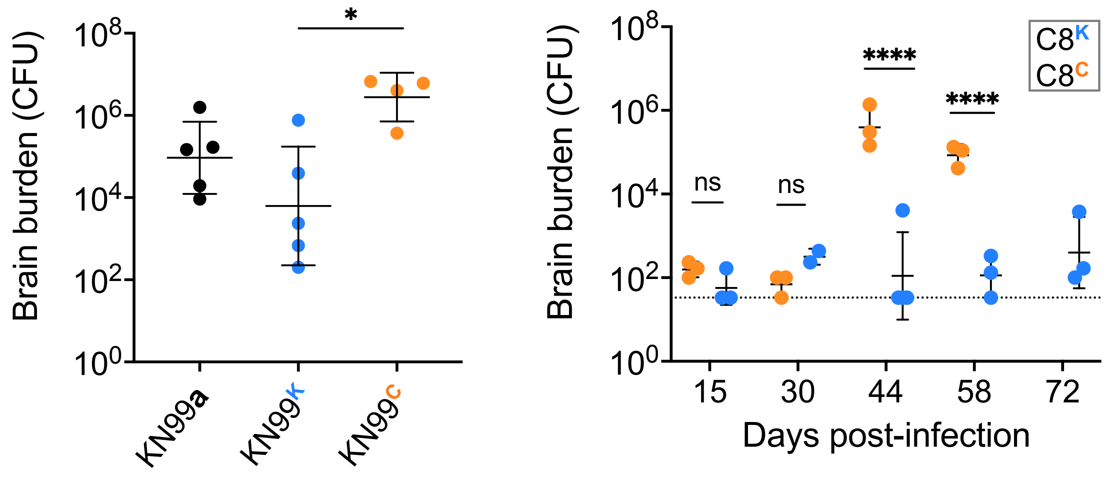


**Supplementary Figure 5.** Brain burden for reciprocal swaps of CKF44_07470 in otherwise identical strains shows that the C8 allele (orange) increases brain CFU of KN99 (left) and the KN99 allele (blue) decreases the brain burden of C8 (right). Mean ± SD of lung burden is shown at 9 days (left) or the indicated times (right); each symbol represents one mouse. P-values were computed by t-test: *, $p < 0.05$; **** $p \leq 0.0001$. Mice infected with C8$^{CCC}$ succumbed to infection between days 58 and 72, so there are no CFU results for these mice at the last time point. Infection was longer for C8 infections because of its baseline low virulence. The parental KN99**a** strain (black) is shown for reference only; it cannot be directly compared to the engineered strains. Dotted line, limit of detection.

**SUPPLEMENTARY FIGURE 6**

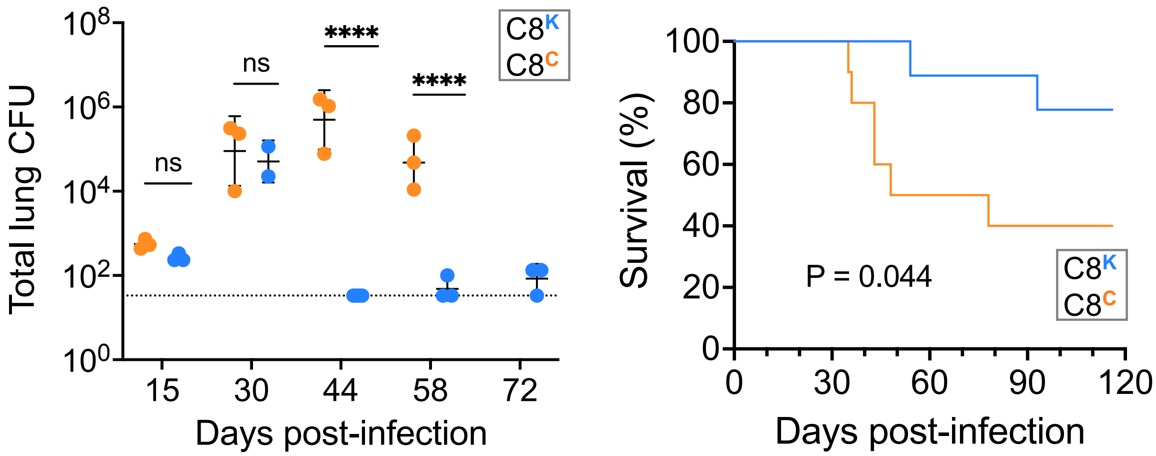


**Supplementary Figure 6.** Sequence swaps show that the KN99 allele of CKF44_07470 decreases the virulence of C8. Strain designations are as above: the base name indicates the background strain and the superscript indicates the allele that was swapped into that background. *Left*, mean ± SD of lung burden at the indicated times after mouse infection; each symbol represents one mouse. Mice infected with $C8^{C}$ (orange symbols) succumbed to infection between days 58 and 72, so there are no CFU results for these mice at the last time point. Dotted line, limit of detection. ****, $P \leq 0.0001$ by t-test. Brain burdens are shown in Supplemental Figure H. *Right*, mouse survival over time, analyzed by Log-rank test. Time courses were extended for C8 infections because of its low baseline virulence.
